# Early-life environmental enrichment generates persistent individualized behavior in mice

**DOI:** 10.1101/851907

**Authors:** Sara Zocher, Susan Schilling, Anna N. Grzyb, Vijay S. Adusumilli, Jadna Bogado Lopes, Sandra Günther, Rupert W. Overall, Gerd Kempermann

**Affiliations:** German Center for Neurodegenerative Diseases (DZNE) Dresden, Tatzberg 41, 01307 Dresden, Germany; Center for Regenerative Therapies Dresden (CRTD), Technische Universität Dresden, Fetscherstraße 105, 01307 Dresden, Germany

## Abstract

Individuals differ in their response to environmental stimuli, but the stability of individualized behaviors and their associated changes in brain plasticity are poorly understood. We developed a novel model of enriched environment to longitudinally monitor 40 inbred mice exploring 35 connected cages over periods of three to six months. We show that behavioral individuality that emerged during the first three months of environmental enrichment persisted when mice were withdrawn from the enriched environment for three additional months. Behavioral trajectories were associated with stable inter-individual differences in adult hippocampal neurogenesis and persistent epigenetic effects on neuronal plasticity genes in the hippocampus. Using genome-wide DNA methylation sequencing, we show that one third of the DNA methylation changes were maintained after withdrawal from the enriched environment. Our results suggest that, even under the most constraint conditions controlling genes and environment, early-life experiences result in lasting individualized changes in behavior and brain plasticity.

## Introduction

Behavioral variability is widespread among species, including humans^1^, and has been attributed to gene-environment interactions and developmental stochasticity^2–4^. In humans, there is high variability in cognitive aging trajectories^5^, which differ between individuals in terms of rate and severity of the age-related cognitive decline. To understand sources of individualized aging, the concepts of “brain reserve” and “brain maintenance” have been introduced^5,6^. While “brain reserve” refers to the accumulation of neural resources during early life that attenuate later functional decline during aging, “brain maintenance” describes the preservation of brain integrity over time. Cognitive and sensory stimulation, particularly early in life, have been linked to increased brain health and resilience against disease^7,8^. We postulate that variability in responses to early-life environmental stimulation could lead to stable brain individualization which serves as a potential substrate for individual “brain reserve”.

We have previously shown that environmental enrichment (ENR) promotes the development of behavioral variability between inbred C57BL6/JRj mice and proposed that ENR could be used as a tool to study brain individualization^9–11^. In those experiments, inter-individual differences among inbred mice emerged when they were housed in the same ENR cage, i.e. even in the absence of nominal genetic and environmental variation. The complexity of ENR allows the individual experience of non-shared environmental components, which amplify and reinforce subtle initial differences between animals over time^12^. However, since these studies were restricted to an ENR housing period of three months, the within-animal stability of the individual ENR-induced differences over longer periods of time and their plasticity towards environmental change remained unelucidated. Would prolonged ENR lead to even further phenotypic divergence or does the individualizing effect plateau?

Activity-dependent brain plasticity is a neurobiological basis for stable behavioral changes that develop in response to environmental stimulation^13^. ENR leads to structural changes in the brain that correlate with increased synaptic plasticity and improved cognitive performance^14–16^. A prime example of ENR-induced structural brain plasticity is the greater number of new neurons that are generated in the hippocampus of ENR-housed mice^17^. Our previous study has shown that, among a number of measures related to structural brain changes, adult hippocampal neurogenesis was the only phenotype, which, in ENR animals, showed enhanced variance in addition to increases in group means that have been reported in essentially all previous studies^11^. Furthermore, we have found that a substantial part of the variance in adult hippocampal neurogenesis (22 %) was explained by individualized behavioral trajectories that emerged during time in ENR^9^, linking individual experience with brain plasticity. Moreover, one of the earliest studies using ENR has shown that increases in cerebral cortex weight in ENR-housed rats were stable for several weeks after withdrawal from ENR^18^, suggesting that ENR-stimulated effects on brain plasticity are potentially long-lasting. This raised the question whether ENR-induced inter-individual differences in adult hippocampal neurogenesis and correlated behaviors are stable and maintained after environmental change.

To analyze the stability of individuality in ENR, we designed a novel cage system that allows the automated longitudinal monitoring of laboratory mice in a large enriched environment over longer time periods. The established system offers greatly improved temporo-spatial resolution of animal activity compared to the enclosure which we had used to establish the emergence of individuality in our previous study^9^. We now asked how behavioral and structural brain individualization would be maintained either upon prolonged or discontinued stimulation. Thereby our study was a direct biological implementation of the concepts of “brain maintenance” and “brain reserve”.

## Results

### Longitudinal monitoring of mice in a stimulus-rich environment

To investigate the effects of long-term ENR on behavioral variability, groups of 40 female C57BL/6JRj mice were housed in ENR or standard housing (STD) for a period of six months (Fig. 1a). To analyze the maintenance of individual behaviors after stimulus withdrawal, a third equally sized animal group was housed in ENR for three months (experimental phase 1) and afterwards returned to STD cages where the animals stayed for another three months (experimental phase 2; ENR-STD mice).

**Fig. 1:**
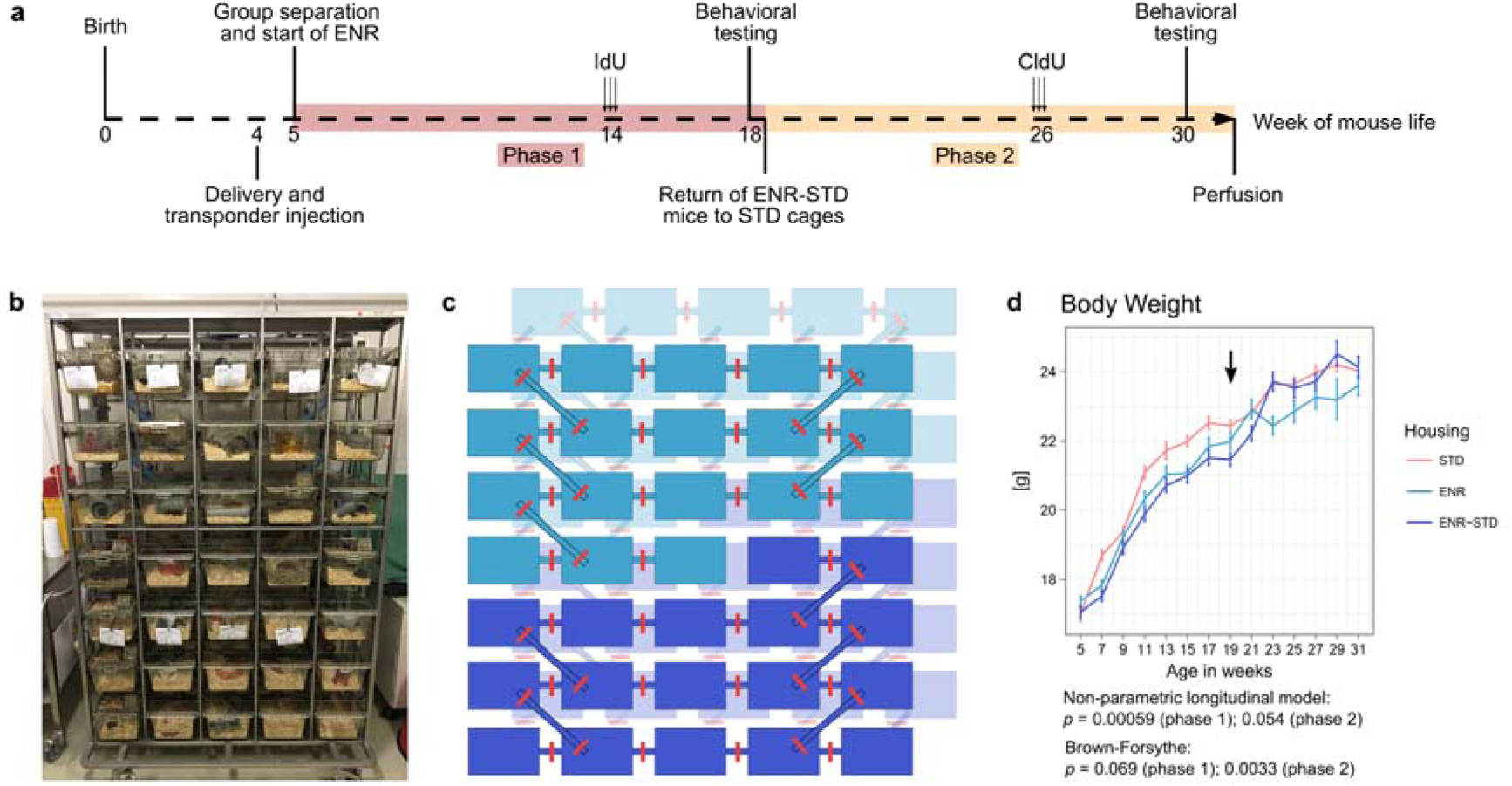
Experimental setup, enriched environment and longitudinal monitoring of body weight. **a,** At an age of five weeks, 120 female C57BL/6JRj mice were split into three equally sized groups: 40 mice lived in environmental enrichment (ENR) for six months, while a second group stayed in standard housing cages (STD) for six months and the third group (ENR-STD) lived in ENR for the first three months (phase 1) and in STD for the last three months of the experiment (phase 2). Behavioral testing was performed with all animals in the last weeks of both phases. For quantification of adult hippocampal neurogenesis, all mice were injected with the thymidine analogues 5-iododeoxyuridine (IdU) and 5-chlorodeoxyuridine (CldU) four weeks before the ends of phase 1 and phase 2, respectively. **b,** Image of the ENR cage system used for automated behavioral tracking of mice by radio frequency identification (RFID) technology. **c,** Schematic representation of the ENR cage depicting cage and tunnel access for the ENR group (light blue; top) and ENR-STD group (violet; bottom). RFID antennae were located around every tunnel and are highlighted in red. **d,** ENR-induced reduction in body weight rebound to STD levels after returning ENR-STD mice to STD cages. Data points indicate means and standard error of the means. Arrow marks return of ENR-STD to STD cages. Depicted *p*-values correspond to housing effects from statistical tests. Full information on statistical tests is presented in Table 1 and Supplementary data 1.

**Table 1:**
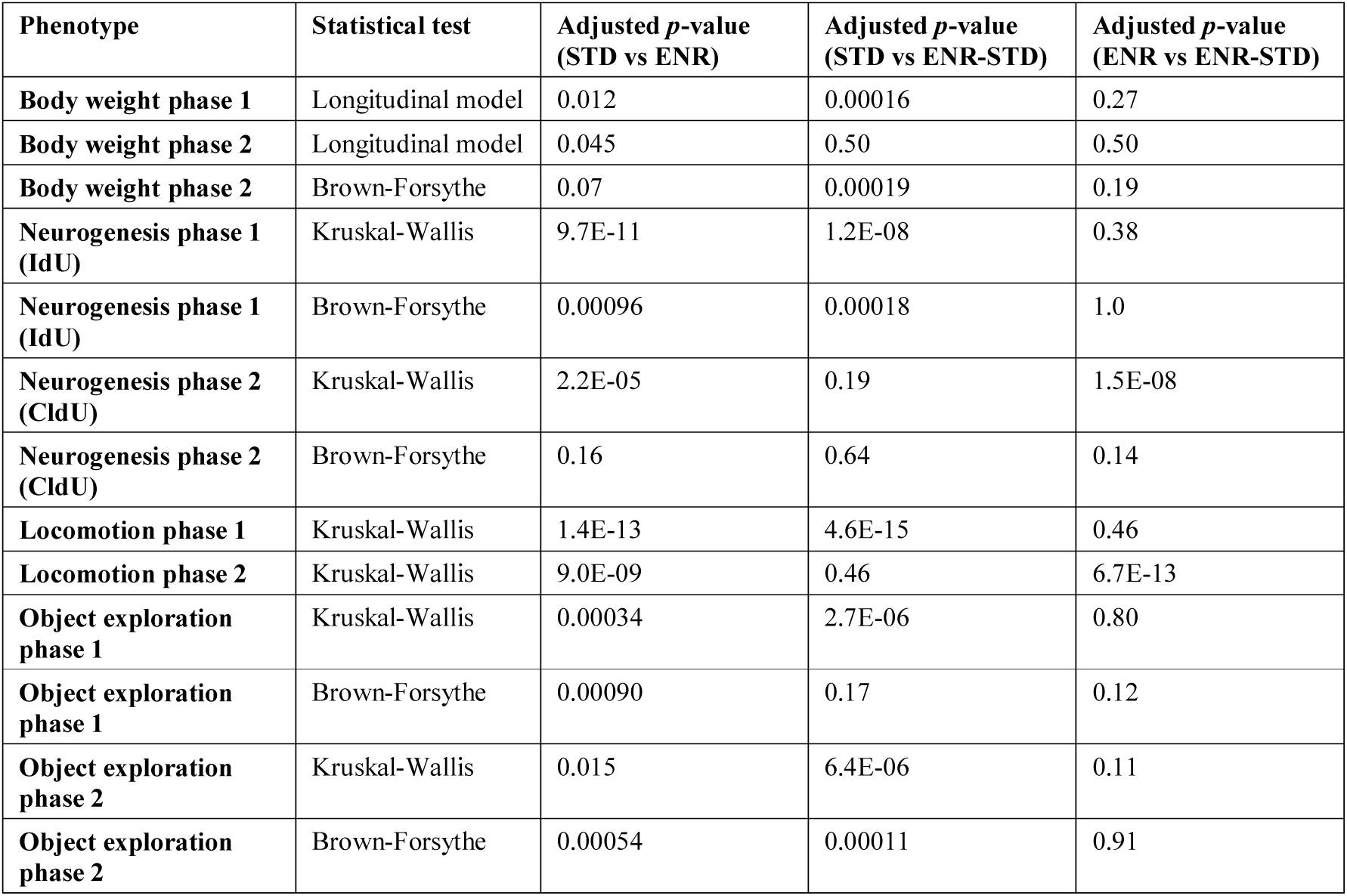
Summary statistics for individual group comparisons

To longitudinally monitor the behavior of indiviudal mice, we used a novel ENR cage system that enabled automated behavioral tracking of animals in a semi-naturalistic environment (Fig. 1b). The ENR cage consisted of 70 interconnected cages arranged on seven levels. Cages were connected by tunnels equipped with antennae for radio frequency identification (RFID)-based detection of animal movements between adjacent cages or levels. During experimental phase 1, ENR and ENR-STD mice stayed in separated sub-compartments of the ENR system, each consisting of 35 cages that covered a total area of 1.37 m^2^ (Fig. 1c). ENR mice remained in the same sub-compartment during experimental phase 2.

To monitor the general health status of mice, we measured body weight of all mice throughout the experiment and found an expected age-related increase in all animal groups (Fig. 1d). At the end of phase 1, body weight of ENR and ENR-STD mice was reduced compared to STD mice (Fig. 1d; Table 1), which is consistent with the previously reported decrease of body weight by ENR^11,17^. However, within four weeks after returning ENR-STD mice to STD, their body weights increased to the levels observed in STD mice, while the long-term ENR group maintained lower body weights throughout phase 2 (Fig. 1d). This result showed that ENR-induced changes in body weight as a gross effect of the stimulation did not persist for long-term after environmental change.

### ENR mice develop stable individualized trajectories in exploratory behavior

We have previously shown that behavioral individualization in ENR can be detected using roaming entropy (RE) as a measure of territorial coverage and spatial exploration of the environment by individual mice^9,10^. Using the longitudinal mouse activity data obtained from the RFID recordings in the ENR cage, we calculated REs for the 40 mice of the ENR group and analyzed the development and stability of individual behavior during the six months of ENR housing.

Mean REs of the mice decreased with time in ENR (Fig. 2a), presumably as a result of the habituation of mice to the ENR cage. Despite this decline in exploratory activity, total variances of nightly REs increased during ENR housing (Fig. 2b), which was accompanied by the divergence of the behavioral trajectories of the individual mice with time (Fig. 2c). To assess the consistency of individual behavior over time, we estimated the variance fraction that is explained by inter-individual behavioral differences and calculated the repeatability of REs within time blocks of 21 calendar days. Repeatability is a measure used in behavioral ecology to detect stable differences between animals in a population that are high relative to intra-individual fluctuations^19,20^. We found that during the first week in ENR (time block T1), the behavior of mice was characterized by repeatability and inter-individual variance not significantly different from zero (Fig. 2d, e; Supplementary table 1). After this initial phase, repeatability of behavior and inter-individual variance continuously increased until the fourth month of ENR housing (time block T5), suggesting a progressive behavioral individualization of mice in ENR. Accordingly, the model assuming heterogeneous inter-individual variance of REs between time blocks was better supported than the model with homogeneous variance (Δdeviance information criteria = 36.8; Supplementary table 2). Thereafter, repeatability did not further increase, resulting in a stabilization of behavioral trajectories, which was supported by the high inter-individual correlations of REs between time blocks in later periods of ENR housing (Fig. 2f; Supplementary Table 3).

**Fig. 2:**
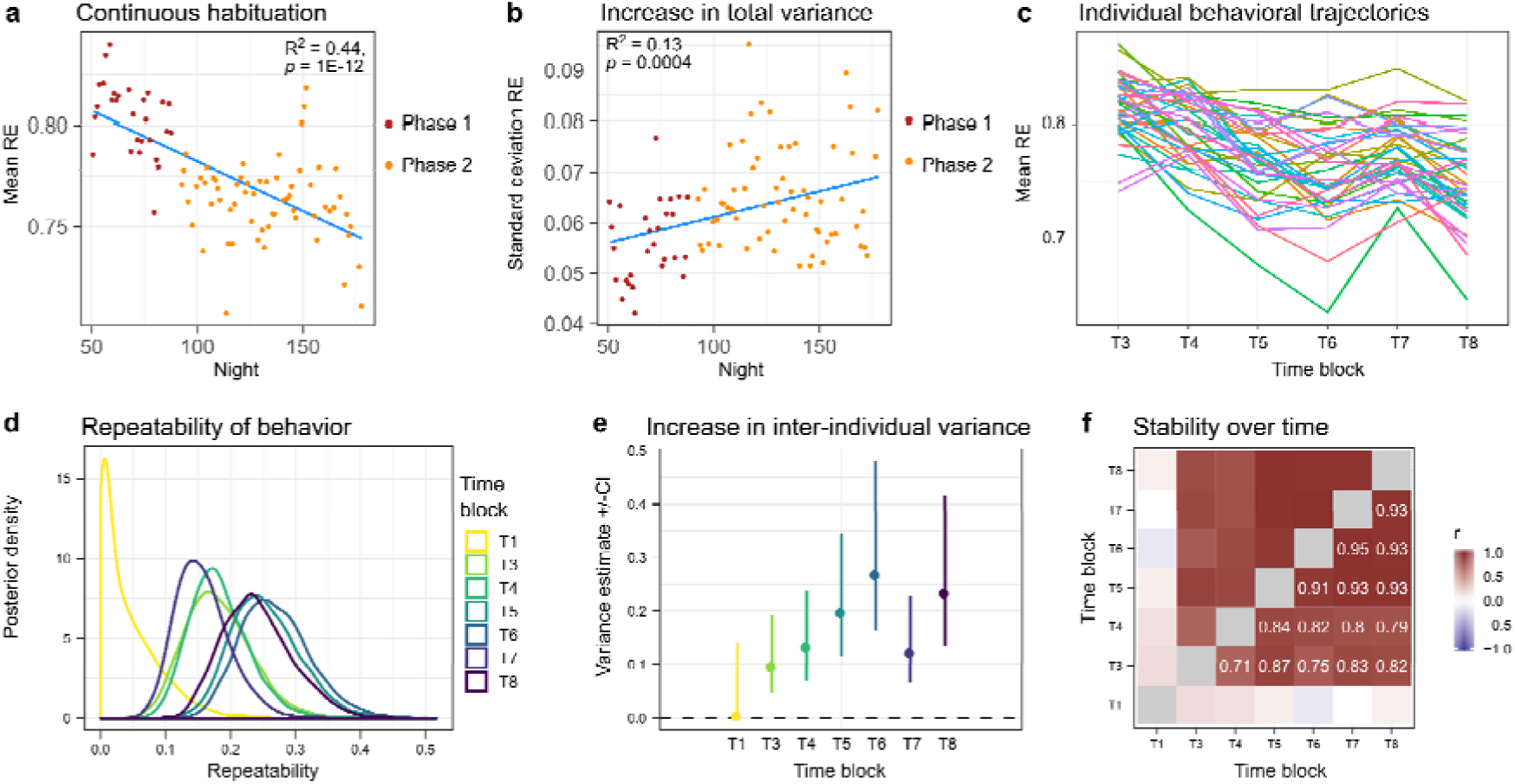
Mice in ENR develop consistent inter-individual differences in behavior. **a,** Mean nightly roaming entropy (RE) of all ENR mice decreased with time. Data prior night 50 was incomplete as a result of technical problems and excluded from the analysis. **b,** Standard deviation of nightly REs increased with time. R^2^ and *p*- values in a-b are derived from linear regression. **c,** Individual mouse trajectories of REs aggregated over time blocks of 21 calendar days diverged with time. **d,** Repeatability of REs between time blocks increased until T5 (fourth month of ENR housing) and remained stable thereafter. Posterior densities depict probabilities of the variance values obtained from generalized linear mixed model. **e,** Variance due to inter-individual behavioral differences increased until T6. Shown are modes of the posterior density with 95 % credible intervals (CI). **f,** Inter-individual correlations of RE were significant between later time blocks. Color code highlights the coefficient of Pearson correlation. See Supplementary tables 1-3 for statistical details.

These data reproduce our previously observed development of behavioral individuality in ENR^9^ in a novel ENR cage system and additionally demonstrate that behavioral trajectories get progressively individualized until the fourth month in ENR and stabilize thereafter.

### Stability of inter-individual differences in adult neurogenesis with time

ENR enhances variance in adult hippocampal neurogenesis compared to STD mice, which we have previously related to the development of stable behavioral trajectories^9,11^. To analyze whether ENR has lasting consequences on individual rates of adult neurogenesis, we injected mice four weeks before the end of each phase with the thymidine analogues 5- iododeoxyuridine (IdU; phase 1) and 5-chlorodeoxyuridine (CldU; phase 2) which can be separately detected by immunohistochemistry (Fig. 1a). ENR and ENR-STD mice showed a greater than 2-fold increase in the numbers of IdU-positive cells in the dentate gyrus and a significantly higher variance of cell numbers compared to STD mice (Fig. 3a; Table 1). Together with the previously observed increase in mean and variance of adult neurogenesis after three months of ENR^11^, these results indicated that individual differences in numbers of neurons generated in the hippocampus during early-life ENR are stable with time and maintained even after withdrawal of ENR.

**Fig. 3:**
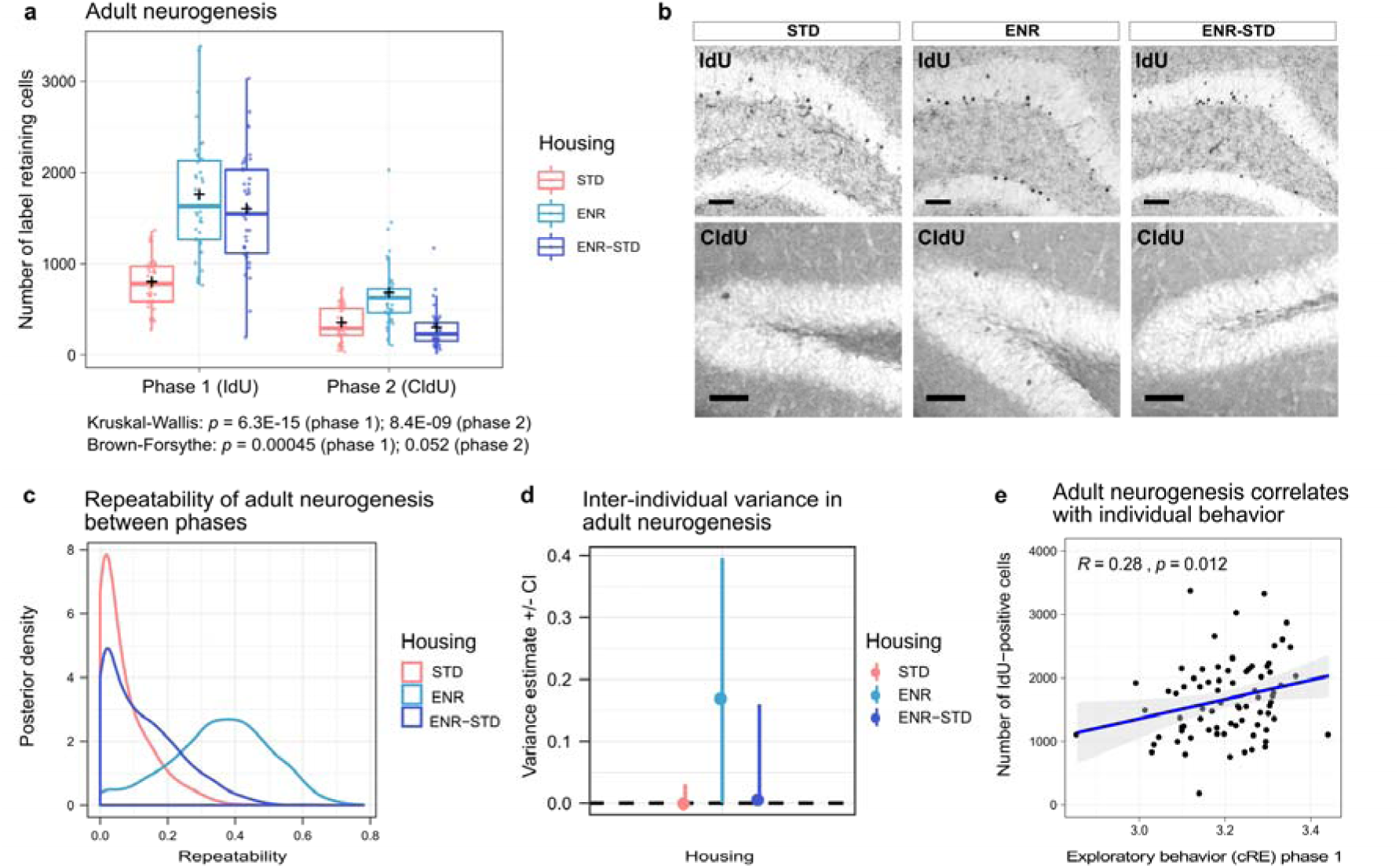
Long-term maintenance of inter-individual differences in adult-born neurons in the hippocampus. Adult neurogenesis during phase 1 and phase 2 was analyzed by injection of IdU (four weeks before the end of phase 1) and CldU (four weeks before the end of phase 2). **a,** Increased mean and variance in numbers of new hippocampal neurons produced during phase 1 (IdU) were observed in ENR and ENR-STD mice at the end of phase 2. ENR but not ENR-STD mice showed increased levels of adult neurogenesis compared to STD mice in phase 2 (CldU). Box and whisker plots: center line - median; plus sign - mean; upper and lower hinges - first and third quartiles; whiskers - highest and lowest values within 1.5 times the interquartile range outside hinges; dots - individual data points. **b,** Representative images of immunohistochemical stainings for detection of IdU- and CldU- positive cells. Scale bar– 50 µm. **c,** Adult hippocampal neurogenesis is repeatable between phases only in ENR mice. **d,** Variance fraction that is explained by inter-individual differences shows posterior distribution shifted towards positive values in ENR compared to STD and ENR-STD mice. Shown are modes of posterior densities with 95 % credible intervals. **e,** Phenotypic correlation of adult neurogenesis to cumulative roaming entropy (cRE) in phase 1. See Supplementary table 1 and Supplementary data 1 for statistical details.

When we compared the numbers of new hippocampal cells generated in both phases, we found an expected age-related decrease in adult hippocampal neurogenesis in all animal groups (Kruskal-Wallis test with Dunn’s posthoc test; STD: *p* = 2.0E-04; ENR: *p* = 8.3E-08; ENR-STD: *p* = 1.1E-18). ENR-STD mice exhibited a lower number of cells generated during experimental phase 2 (CldU-positive) compared to ENR mice but no difference compared to STD mice (Fig. 3a; Table 1), indicating that ENR-stimulated neurogenic activity is not maintained after withdrawal of environmental stimulation. In contrast, ENR mice showed a significant 2-fold increase in CldU-positive cell numbers in the dentate gyrus compared to STD mice. Additionally, numbers of new hippocampal cells were repeatable and significantly correlated between phases only in ENR mice (Fig. 3c; Supplementary Fig. 1), which suggested that ENR mice developed consistent inter-individual differences in adult neurogenesis. Although the lower 95 % confidence intervals for inter-individual variance in ENR mice abutted zero (Fig. 3d), the model which estimated separate inter-individual variance in housing groups was better supported than a model with homogeneous variance (Δdeviance information criteria = 13.6; Supplementary table 2). Note that the survival periods of IdU- and CldU-labeled cells were different (four and one months, respectively). Our results also indicate that despite the discontinuation of the environmental stimulation in the ENR- STD group, the new neurons generated in response to the initial ENR phase were maintained, while the *de novo* production of new neurons declined to baseline levels.

To relate adult hippocampal neurogenesis to the emerged behavioral trajectories, we correlated exploration in ENR (cumulative roaming entropy) with adult neurogenesis (number of IdU-positive cells) for all mice that were housed in ENR during phase 1 (ENR and ENR- STD group). In agreement with our previous observations^9^, adult neurogenesis showed a positive correlation with individual levels of exploratory behavior (Fig. 3f).

Taken together, these results showed that (1) individual behavioral trajectories and correlated levels of adult hippocampal neurogenesis developed in ENR during phase 1; (2) individuality in behavior and adult neurogenesis remained stable with continued ENR in phase 2; and (3) structural individuality of the hippocampus through adult neurogenesis in phase 1 was maintained after withdrawal of ENR in phase 2.

### Selective maintenance of ENR-induced individuality in object exploration after environmental change

We next asked whether the stability of the emerged individual behaviors with time was dependent on continuous stimulation in ENR or whether behavioral patterns were also maintained after returning mice to STD cages as suggested by the lasting individualized hippocampal structures. We have recently shown that ENR-induced behavioral variability can be detected using open field and object exploration tests in a cross-sectional experimental design^11^. To investigate the long-lasting effects of ENR on behavioral variability, all mice were analyzed in open field and object exploration tests at the end of both experimental phases (Fig. 1a).

ENR changed activity patterns in the open field in phase 1, which were preserved with prolonged time in ENR but not maintained after ENR-STD mice returned to STD cages in phase 2 (Supplementary Fig. 2). For instance, locomotor activity in the open field arena was reduced in ENR mice compared to STD mice at the end of phase 1 and phase 2 (Fig. 4a-b, Table 1). In contrast, while locomotion of ENR-STD mice was reduced compared to STD mice in phase 1, it was similar to STD mice and increased compared to ENR mice in phase 2. Comparison of locomotion between phase 1 and phase 2 further showed that individual locomotor activity was stable within animals in STD and ENR mice, but this stability was lost in ENR-STD mice (Fig. 4c-d; Supplementary Fig. 3).

**Fig. 4:**
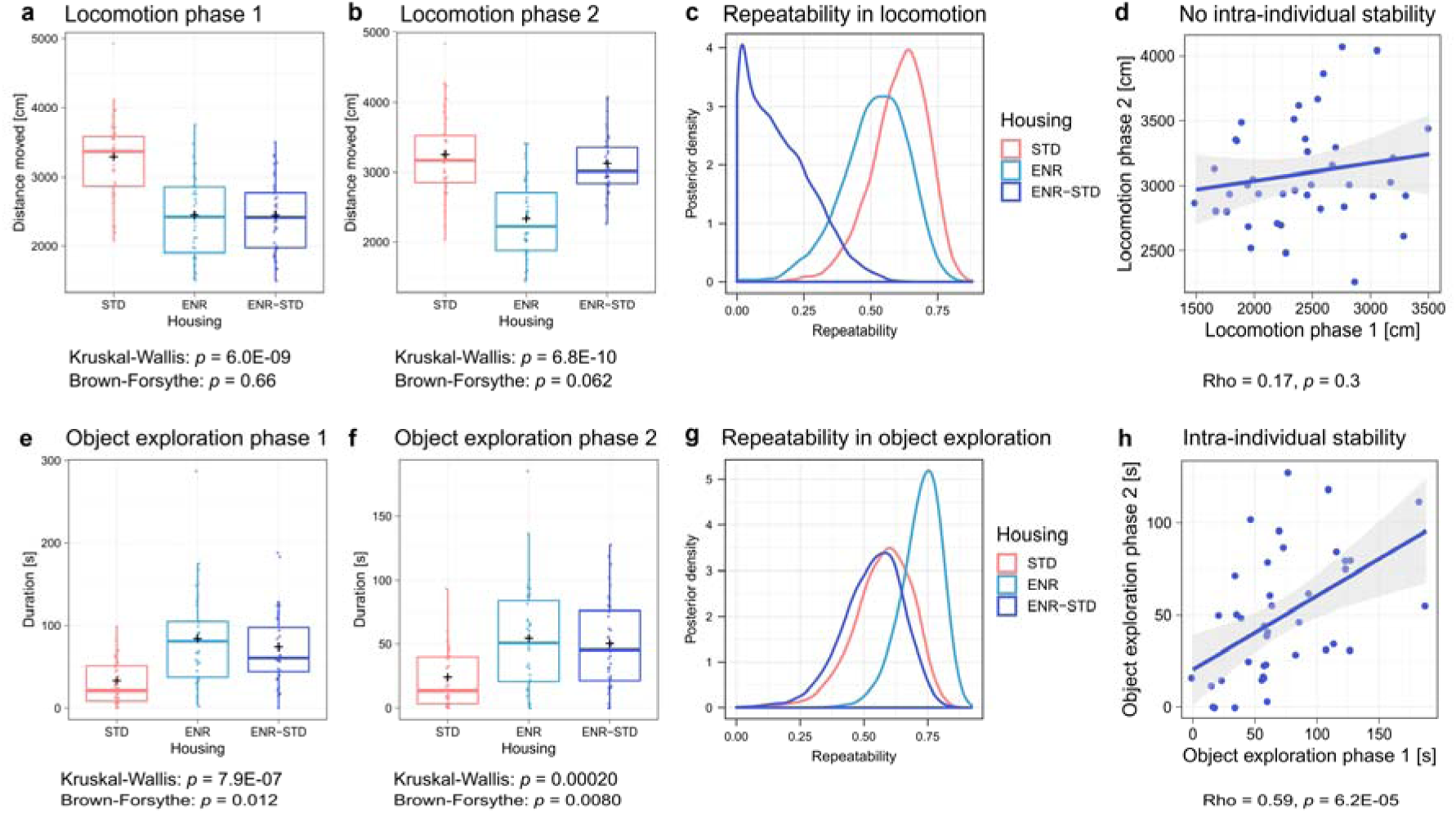
Stability of ENR-induced individuality in object exploration with time and maintenance after withdrawal from ENR. **a-d,** Locomotion in the open field test is depicted as s an example of a behavior that is not maintained after withdrawal from ENR. **a,** At the end of phase 1, ENR and ENR-STD mice traveled shorter distances in the open field test compared to STD mice. **b,** At the end of phase 2, ENR mice maintained reduced levels of locomotion, while ENR-STD mice traveled similar distances as STD mice. **c,** High repeatability of locomotion between phase 1 and phase 2 is observed in ENR and STD mice but not in ENR-STD mice. **d,** Lack of correlation of individual locomotion between phase 1 and phase 2 in ENR-STD mice suggests behavioral change after withdrawal from ENR. **e,** ENR increased duration and variance of object exploration in trial 1 of the object exploration test. **f,** Increases in mean and variance of initial object exploration induced by ENR can still be observed after withdrawal from ENR. **g,** High repeatability of initial object exploration between phase 1 and phase 2 in ENR and ENR-STD mice indicates maintenance of individual levels of object exploration after withdrawal of ENR. **h,** Significant correlation of initial object exploration between phases in ENR-STD mice confirms intra-individual stability of behavior. Box and whisker plots in a, b, e, f as described in Fig. 3. See Supplementary table 1 and Supplementary data 1 for statistical details.

In contrast to the animals’ activity in the open field test, we found that the ENR-induced variability in object exploration was not only stable over time in ENR but also maintained after withdrawal from ENR. In the object exploration tests of phase 1 and 2, ENR mice spent more time around objects and showed a significantly higher variance of the duration of object exploration compared to STD mice (Fig. 4e-f; Table 1; Supplementary Fig. 2). This result confirms the previously observed variability in object exploration after three months of ENR^11^ and additionally demonstrates that the variance-enhancing effect of ENR in object exploration is preserved with prolonged periods of ENR housing. Remarkably, in both phases ENR-STD mice showed greater means and variances of object exploration compared to STD mice and no differences compared to ENR mice, suggesting that ENR-induced changes in object exploration are maintained after returning mice to STD cages. Moreover, individual levels of initial object exploration were highly repeatable and significantly correlated between the two experimental phases not only in STD and ENR mice but also in the ENR-STD group (Fig. 4g-h; Supplementary Fig. 3). These results indicate that ENR-induced increases in object exploration are stably maintained within animals during time in ENR and even after withdrawal of animals from the enriched environment.

In summary, we showed that ENR-induced individualization in behavior and adult hippocampal neurogenesis in phase 1 resulted in lasting behavioral and structural brain individuality that was not only preserved with continuous ENR housing in phase 2, but also maintained for three months after withdrawal from ENR.

### Maintenance of ENR-induced DNA methylation changes in the dentate gyrus after environmental change

Experience-dependent epigenetic changes, such as DNA methylation, have been linked to stable behavioral differences^21^. Dynamic DNA methylation changes in neurons contribute to synaptic plasticity and memory formation^22,23^. To identify molecular mechanisms underlying the maintenance of individual behavior after withdrawal of ENR, we performed genome-wide DNA methylation profiling on micro-dissected dentate gyrus tissue by reduced representation bisulfite sequencing^24,25^.

We detected significant methylation differences between ENR and STD mice at 12,167 CpGs (2.67 % of all CpGs) and 1,927 CpHs (0.087 % of CpHs; Supplementary Fig. 4a-b; Supplemental data 2). Genes containing ENR-induced differentially methylated cytosines were enriched in pathways related to axon guidance and neuronal plasticity (Supplementary Fig. 4c-d). Comparing ENR-STD with STD mice, we identified 10,216 differentially methylated CpGs (2.24 % of CpGs) and 1,315 differentially methylated CpHs (0.053 % of CpHs; Supplementary Fig. 4e-f; Supplemental data 2), indicating that early-life ENR led to stable DNA methylation changes in the dentate gyrus that persisted after withdrawal of the stimulus. Genes with differentially methylated cytosines after ENR-STD showed a similar enrichment in axon guidance and neuronal plasticity pathways as long-term ENR mice (Supplementary Fig. 4g-h).

To evaluate whether DNA methylation patterns of ENR-STD mice resemble those in ENR animals, we first correlated the methylation differences of ENR and ENR-STD compared to STD for all individual cytosines. Significant positive correlations were detected for CpG (*r* = 0.55) and CpH (*r* = 0.55) contexts (Fig. 5a), suggesting that ENR and ENR-STD showed similar methylation changes at individual cytosines on a genome-wide scale. Next, we overlapped significantly differentially methylated cytosines by ENR with cytosines differentially methylated by ENR-STD (Fig. 5b). In total, 27.30 % of ENR-induced differentially methylated CpGs and 31.40 % of CpHs also showed similar methylation changes after ENR-STD. Plotting the absolute DNA methylation percentages for those overlapping cytosines showed that the magnitude of the methylation changes was similar in ENR and ENR-STD mice (Fig. 5c). While STD mice showed accumulations of CpGs and CpHs with intermediate methylation percentages, ENR led to long-lasting hypermethylation of those CpGs and a hypomethylation of the CpHs with intermediate methylation percentages (Fig. 5c). These results suggested that ENR led to DNA methylation changes that are maintained for at least three months after withdrawal from ENR.

**Fig. 5:**
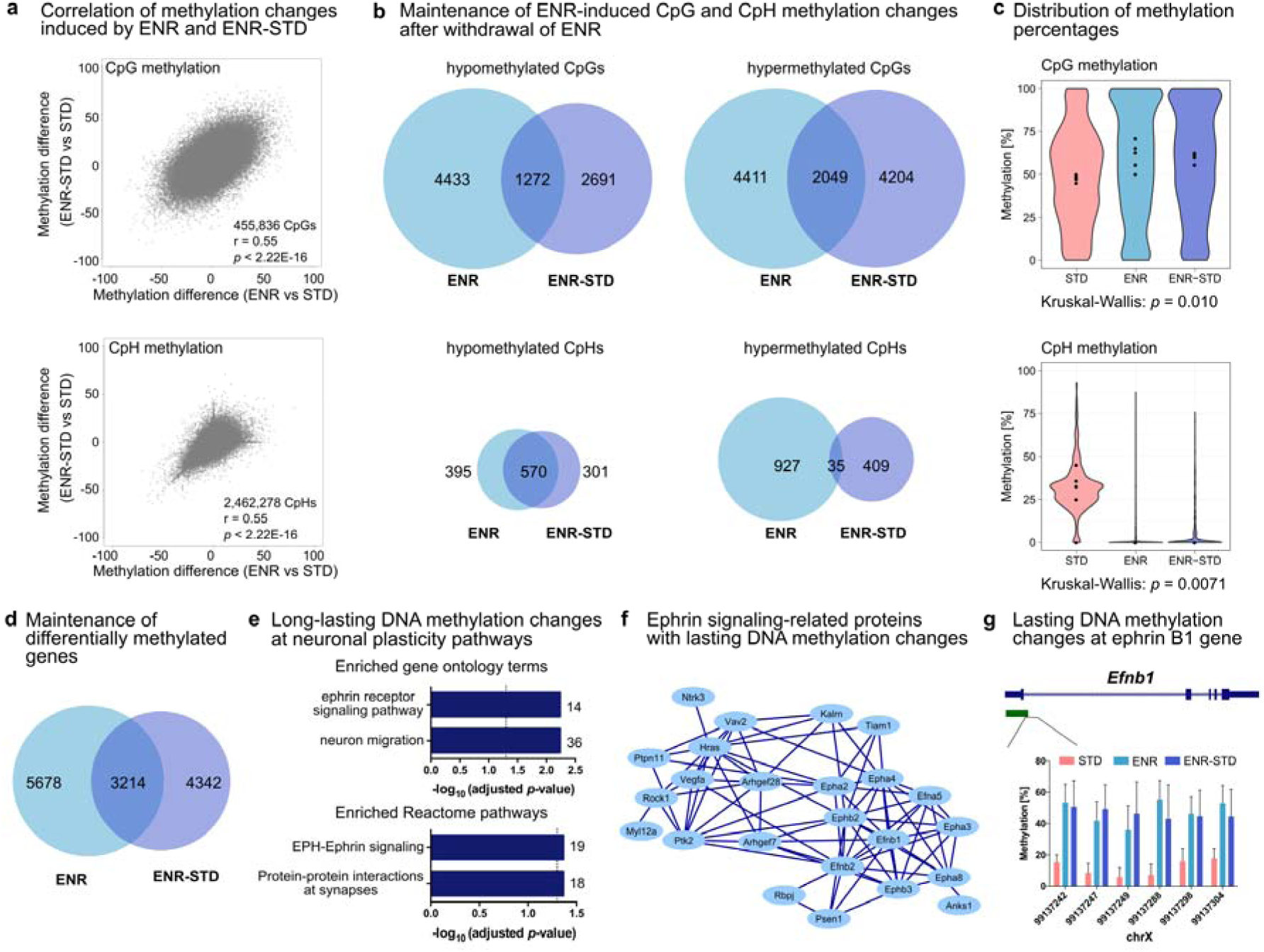
ENR led to long-lasting DNA methylation changes in the dentate gyrus at ephrin signaling-related genes. **a,** Differences in CpG and CpH methylation between ENR and STD mice correlate with methylation differences between ENR-STD and STD mice at individual cytosines (all cytosines sequenced). *P*-values from Pearson correlation (n = 5 per group). **b,** Overlap of 27.30 % of differentially methylated CpGs and 31.40 % of differentially methylated CpHs between ENR-induced and ENR-STD-induced methylation changes. **c,** ENR- STD mice show absolute DNA methylation percentages of the overlapping 3,321 CpGs and 605 CpHs similar to ENR mice but different to STD mice. Median methylation percentages for individual samples are highlighted. **d,** Overlap of genes with differentially methylated CpGs or CpHs in ENR compared to STD and ENR-STD compared to STD mice. **e,** The 3,214 genes with long-lasting ENR-induced DNA methylation changes are enriched in biological processes (gene ontology) and pathways (Reactome) related to neuronal plasticity, including ephrin signaling, neuron migration and synaptic plasticity pathways. **f,** STRING interaction network of genes with enrichment in ephrin receptor signaling pathways. **g**, ENR-induced DNA methylation changes at CpGs located in proximity to exon 1 of the *Efnb1* gene are maintained in ENR-STD mice. Green bar depicts a CpG island. Bars represent mean and standard error of the mean for each CpG.

Interestingly, context-specific differences were detected in the directionality of the persistent methylation changes. While in the CpG context, 22.30 % and 31.72 % of ENR-induced differentially methylated cytosines also changed in ENR-STD mice, in the CpH context, 59.07 % of hypomethylated CpHs but only 3.64 % of hypermethylated CpHs were maintained (Fig. 5b). This indicates that, at CpHs, maintenance of ENR effects is restricted to hypomethlyation and that ENR-induced CpH hypermethylation is dependent on continuous stimulation in ENR and does not persist after withdrawal of ENR.

Gene annotation showed that 36.14 % of ENR-induced differentially methylated genes persisted in ENR-STD mice (Fig. 5d). Gene ontology and Reactome pathway analysis suggested that genes with maintained methylation changes were involved in processes related to ephrin signaling, neuron migration and synaptic signaling (Fig. 5e). Notably, 28 genes with known role in ephrin signaling showed persistent ENR-induced DNA methylation changes (Fig. 5f). One example of these was a differentially methylated region in proximity to the transcription start site of the ephrin B1 ligand *Efnb1* (Fig. 5g). Ephrin signaling has multiple roles in neuronal plasticity such as axon guidance, control of dendritic growth, synapse formation, synaptic plasticity^26^, neuronal differentiation and adult hippocampal neurogenesis^27^. Furthermore, ephrin signaling controls behaviors related to anxiety, stress and depression^28–30^. Hence, the ENR-induced regulation of the ephrin signaling pathway in the hippocampus is a potential molecular link between persistent individual behavior and brain plasticity after withdrawal of ENR.

## Discussion

A characteristic feature of behavioral individuality is the stability of behavioral patterns over time^31,32^. Here, we showed that variability in behavior and adult hippocampal neurogenesis induced by ENR are stable within animals over prolonged periods of ENR housing. Furthermore, structural individualization of the hippocampus through long-term integrated adult-born neurons was maintained even after withdrawal of ENR, together with sustained individualized behavior in object exploration. Persistent behavioral differences after ENR were accompanied by changes in hippocampal DNA methylation patterns, which were maintained after stimulus withdrawal. Our data reveal the long-term maintenance of behavioral individuality in ENR and its relation to structural plasticity and the epigenetic state of the hippocampus.

ENR increased inter-individual differences in spatial exploration (RE) and variance in object exploration and adult hippocampal neurogenesis, which confirmed the findings from our previous studies^9,11^. The fact that we detected ENR-induced inter-individual differences in all three studies despite using structurally distinct enriched environment cages, highlights the robustness of the development of individuality in ENR. While both previous studies used wide span cages that allowed exploration and interaction in an open space, the current study employed a novel compartmentalized ENR system in which mice can navigate through a network of separated cages. Moreover, in the present study, the two groups ENR and ENR- STD were similarly housed in ENR during experimental phase 1 and tested in open field and object exploration tests afterwards, which served as an internal replication of the initial ENR effect. Both groups showed increased variance in object exploration and adult neurogenesis compared to STD in phase 1 and no significant behavioral differences were observed between them. The robustness and reproducibility of the individualization in behavior and hippocampal structure in ENR underpins the suitability of enriched environments as a tool to study mechanisms of individuality development.

Behavioral individualization, as measured by increasing inter-individual variance and repeatability in RE, progressed until the fourth month of ENR housing and plateaued thereafter. We have previously proposed that individuality in ENR develops by the amplification of initial differences through a self-reinforced positive feedback loop between behavior and adult hippocampal neurogenesis^9,11^, which was supported by the positive phenotypic correlation between neurogenesis and exploratory activity. Other studies also suggested self-reinforcement as drivers of animal variability^33^. The here detected stability of inter-individual differences in adult hippocampal neurogenesis and the significant repeatability of this phenotype only in ENR animals further support this model. The observed age-related decrease in adult hippocampal neurogenesis as well as the reduction of exploratory activity due to the habituation to the ENR cage might have reduced the strength of the positive feedback loop with time and resulted in the detected plateauing of behavioral individualization. Despite this plateauing effect, behavioral trajectories, once developed, were stable over time in ENR, which was highlighted by the high inter-individual correlations of RE between time blocks and by the high repeatability ENR mice showed in object exploration between phases. Future studies should address whether these behavioral trajectories remain stable throughout the animals’ life and result in individualized brain health during aging.

Our results further show that most activity measures, such as locomotion and frequency of center crossings in the open field test as well as body weight, are not maintained post withdrawal of ENR. Object exploration stood out as the one activity measure where the ENR-induced changes were maintained within mice for at least three months after they were withdrawn from the enriched environment. During ENR housing, the weekly change of object location and type (toys, houses and tunnels) in the cage provides repeated cognitive and novelty stimulation and, thus, represents a core aspect of ENR. Previous studies by others have shown stable changes in hippocampal synaptic plasticity^15^, cerebral cortex weight and cortical acetylcholinesterase activity^18^ after withdrawal from ENR, which could contribute to the observed maintenance of individual behavior. We here demonstrated that, although the pro-neurogenic effect of ENR was depended on continuous stimulation, early-life ENR led to persistent individualization of the hippocampal network through adult neurogenesis. Since adult hippocampal neurogenesis has a known role in promoting cognitive flexibility and influencing affective behaviors^34–37^, the lasting behavioral changes could be mediated (at least in part) by the long-term integration of adult-born neurons generated during early-life ENR. Future experiments should investigate the causal role of adult hippocampal neurogenesis and other brain plasticity measures in the development and maintenance of individual behavioral trajectories.

A number of studies suggested epigenetic factors as potential molecular drivers of individuality development^3,33^. DNA methylation profiling in humans identified tissue-independent, inter-individual DNA methylation differences, many of which were attributed to environmental variation, including non-shared environmental influences, rather than genetic differences^38,39^. We have profiled DNA methylation changes that might be associated with the persistence of individuality in exploratory behavior after environmental change. Remarkably, we found that almost one third of the ENR-induced DNA methylation changes in the hippocampal dentate gyrus were maintained for at least three months after returning mice to STD cages. Future investigations into the functional role of the detected ENR-induced DNA methylation changes in the development and maintenance of inter-individual behavioral differences could provide vital links between gene expression and emergent behavioral patterns. Moreover, other molecular players, synergistically with ENR-induced DNA methylation changes or in isolation, can potentiate individualized behavior. For instance, pre-existing inter-individual DNA methylation differences induced by early-life experiences such as maternal care^40^, ENR-induced epigenetic changes other than DNA methylation^41^ or environmentally regulated somatic mosaicism in the hippocampus^42^ are potential drivers of behavioral variability. Furthermore, differential miRNA levels in the hippocampus have been linked to lasting behavioral changes after ENR^15^ and could play a role in the observed maintenance of ENR-induced individuality. Since DNA methylation controls many molecular processes, including expression of miRNAs^43^, somatic mosaicism^44^ and other epigenetic modifications in the brain^45^, the here identified DNA methylation changes likely interact with other molecular players to facilitate the development and maintenance of individual behavior.

Our study suggests that a longitudinal version of the enriched environment paradigm can indeed be applied to study the neurobiological mechanisms underlying fundamental concepts such as “brain reserves” and “brain maintenance”. We show that early exposure to enriched sensory, cognitive and social stimulation induces long lasting changes in behavior, brain plasticity and the hippocampal epigenome and thus highlight the potential of environmental experiences to determine life trajectories. At the same time, we identify plastic responses to ENR that depend on continuous stimulation. Our paradigm thus offers an opportunity to dissect neurobiological mechanisms underlying brain resilience by opening the related animal research to the inclusion of longitudinal trajectories.

## Methods

### Animal husbandry

Female C57BL/6JRj mice were purchased at an age of four weeks from Janvier Labs and housed in groups of five animals in standard polycarbonate cages (Type III; Tecniplast). In the first week upon arrival, all animals were subcutanously injected into their neck with a glass-coated microtransponder (SID 102/A/2; Euro I.D.) under brief isoflurane anesthesia. At an age of five weeks, mice were randomly assigned to three experimental groups using the online tool “Research Randomizer” (www.randomizer.org). Two animal groups of 40 mice each were housed in a cage system custom-built to our specifications (PhenoSys GmbH, now marketed as “PhenoSys ColonyRack Individuality 3.0”), which was divided into two equally sized enriched environments (Fig. 1b,c). Each enriched environment covered an area of 1.37 m^2^ and consisted of 35 polycarbonate cages (1264C Type II, Tecniplast) that were connected via transparent tunnels and distributed on four levels. Food and water was provided on every level of the cage system. For cognitive stimulation, the cages were equipped with plastic toys, tunnels and hideouts which were replaced and rearranged once per week. Additional 40 mice stayed in standard polycarbonate cages (36.5×20.7×14 cm; Type III, Tecniplast) in groups of five animals per cage. Dirty cages in the enriched environment and the standard housing cages were cleaned once per week.

Mice were maintained on a 12 hr light/12 hr dark cycle with 55 ± 10% humidity at the animal facility of the Center for Regenerative Therapies Dresden. Control and enriched animals received the same fortified chow (#V1534; Sniff) with 9 % of energy from fat, 24 % from protein and 67 % from carbohydrates. Food and water were provided freely. All mice received intraperitoneal injections of IdU (57.5 mg/kg) nine weeks after start of the experiment and of CldU (42.5 mg/kg) four weeks before perfusion. IdU and CldU were dissolved in 0.9 % sodium chloride and injected three times at six hour time intervals. All experiments were conducted in accordance with the applicable European and national regulations (Tierschutzgesetz) and were approved by the local authority (Landesdirektion Sachsen; file number 25-5131/354/63).

### Analysis of RFID data

Antenna contacts were recorded using the software *PhenoSoft Control* (PhenoSys GmbH), which saved antenna identifier and mouse identifier together with the time stamp of the antenna contact into a database. During the course of the experiment, over 24.13 mln (phase 1) and 8.52 mln (phase 2) events were registered. Data between 40^th^ and 49^th^ day of ENR were removed from the analysis due to technical problems. Between the 9^th^ and 35^th^ day of ENR, animals repeatedly forced the locks introduced in the tunnels between two compartments of the cage system and thus ENR and ENR-STD group mixed. Because the number and locations at which animal can be recorded influences its distribution and entropy value, the data from this period could not be compared with the rest of the data and were therefore excluded from further analysis.

Data reduction and calculation of roaming entropy (RE) were performed as previously described^9^. Because mice are nocturnal animals, only the events recorded during the dark phase were retained. Each night was divided into 8640 segments of 5 seconds length and for each mouse and each time segment, the last registered antenna was recorded into a time series. Frequencies of antennae in this time series were converted to probabilities *p_i,j,t_*of a mouse *i* being at an antenna *j* at a night *t*. Shannon entropy of the roaming distribution was calculated as: 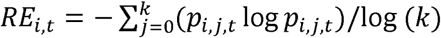; where *k* is the number of antennae. Dividing the entropy by log(*k*) scales the RE to the range between zero and one. Data from nights following events which could disturb patterns of exploration, such as cleaning of the cage or behavioral testing, were excluded. The reduced data set comprised 99 nights, which were partitioned into eight time blocks, each spanning 21 calendar days. The number of nights per each time block in the final data set were as follows: T1: 6; T2: 3; T3: 11; T4: 17; T5: 15; T6: 17; T7: 16; T8: 14. Cumulative RE at the end phase 1 was calculated by cumulative addition of mean RE from the first four time blocks.

### Open field and object exploration tests

The open field and object exploration tests were performed similarly as previously described^11^. Briefly, mice were placed into a square arena (60 × 60 cm) and their exploration was recorded using a camera (Logitech) and EthoVision software (Noldus). In total, four trials were performed on two consecutive days with a trial length of five minutes each (Supplementary Fig. 3a). In the first trial (open field test), mice were put in the empty arena, which also served as habituation for the object exploration tests. In the following two trials, mice were presented with two identical objects. In the fourth trial, one object was replaced with a “new” object. To avoid preference for object properties or placement, the use of object A and object B as “old” or “new” object and the position of the “new” object was randomized. Object A was a composite of a red and yellow plastic tube, each with a diameter of 55 mm and a height of 98 mm. Object B was a grey cuboid of 13 x 10 x 6 cm with a blue surface and holes on one side. Data analysis of open field and object exploration tests was performed as previously described^11^.

### Tissue preparation and immunohistochemistry

Tissue fixation and immunohistochemistry for the analysis of adult neurogenesis were performed as previously described^11^. Briefly, mice were anesthetized with 100 mg/kg ketamine (WDT) and 10 mg/kg xylazin (Serumwerk Bernburg AG) and transcardially perfused with 0.9 % sodium chloride. Brains were removed from the skull and one hemisphere fixed in 4 % paraformaldehyde prepared in phosphate buffer (pH 7.4) overnight at 4 °C. Brains were incubated in 30 % sucrose in phosphate buffer for two days and cut into 40 μm coronal sections using a dry-ice-cooled copper block on a sliding microtome (Leica, SM2000R). Sections were stored at 4 °C in cryoprotectant solution (25 % ethyleneglycol, 25 % glycerol in 0.1 M phosphate buffer, pH 7.4).

For detection of IdU- and CldU-positive cells, the peroxidase method was applied. Briefly, free-floating sections were incubated in 0.6 % hydrogen peroxide for 30 min to inhibit endogenous peroxidase activity. For antigen retrieval, sections were incubated in pre-warmed 2.5 M hydrochloric acid for 30 min at 37 °C, followed by extensive washes. Unspecific binding sites were blocked in Tris buffered saline (TBS) supplemented with 10 % donkey serum (Jackson Immuno Research Labs) and 0.2 % Triton X-100 (Carl Roth) for 1 h at room temperature. Primary antibodies were applied overnight at 4 °C as follows: for IdU detection, monoclonal mouse anti-BrdU (1:500; BD Bioscience) and for CldU detection, monoclonal rat anti-BrdU (1:500, Serotec). Biotinylated secondary antibodies (Jackson Immuno Research Labs) were incubated for 2 h at room temperature. Antibodies were diluted in TBS supplemented with 3 % donkey serum and 0.2 % Triton X-100. Detection was performed using the Vectastain ABC-Elite reagent (9 μg/ml of each component, Vector Laboratories, LINARIS) with diaminobenzidine (0.075 mg/ml; Sigma) and 0.04 % nickel chloride as a chromogen. All washing steps were performed in TBS. Stained sections were mounted onto glass slides, cleared with Neo-Clear (Millipore) and cover-slipped using Neo-Mount (Millipore). IdU- and CldU-positive cells were counted on every sixth section along the entire rostro-caudal axis of the dentate gyrus using a brightfield microscope (Leica DM 750).

### Statistical analysis

Experiments were carried out with the experimenter blind to the experimental group. Statistical analyses were performed using the statistical software R (<u>R Core Team, 2014</u>). To compare means, we used Kruskal-Wallis test with Dunn’s *post hoc* test. To compare variances between groups, Brown-Forsythe test was performed using the *leveneTest* function from the car package. Longitudinal data were analyzed using a rank-based non-parametric test using the *nparLD* function from the nparLD package^46^. All tests were two-tailed and differences were considered to be statistically significant at a *p* < 0.05. Multiple testing correction was performed using Holm method. Data were visualized using the ggplot2 package^47^. In the box-whisker plots, center line and plus sign mark the median and mean, respectively. Upper and lower hinges indicate first and third quartiles. The upper whisker extends from the hinge to the largest value no more than 1.5 times the interquartile range (IQR, a distance between the first and third quartiles); the lower whisker extends from the hinge to the smallest value at most 1.5 times IQR. Full results of statistical tests are available in Supplemental data 1.

### Mixed linear models and repeatability estimation

Repeatability (*R*) is the fraction of total variance which can be attributed to the differences between individuals 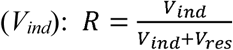; where *V_res_* is a residual, i.e. within individual variance^20,48^. To decompose the phenotypic variance into inter- and within-individual components, we employed generalized linear mixed models in a Bayesian framework implemented in the MCMCglmm package v2.29^49^ in R v3.4.4.

Behavioral phenotypes from open field and object exploration tests were square root transformed to correct the skew and the normal distribution of the data was confirmed using Shapiro-Wilk test. RE and neurogenesis were mean-centered and scaled to unity variance. To calculate repeatability in object exploration between phases, we only used data from the first trial of the object exploration test to avoid effects of habituation. The full model for RE in the ENR animals included the time block as a fixed effect and an interaction between time block and an individual identifier as a random effect, which was specified using the function *us*. This specification of the random term allowed to estimate inter-individual variance for each time block, as well as covariance between the time blocks. The residual variance was estimated separately in each time block using the function *idh*. The models for behavioral parameters and neurogenesis included an interaction between housing and the phase as fixed effects, and an interaction between housing and an individual identifier as a random effect, as well as heterogeneous residual variance specified with the function *idh*. To assess whether inter-individual variance differed between time blocks (RE) or housing conditions (behavior, neurogenesis), we omitted respective terms in the model and compared the deviance information criteria (DIC) between the full and simpler models assuming that a DIC score reduced by at least 2 units indicates better fit to the data.

Models were fitted by Markov chain Monte Carlo estimation with Gibbs sampling using the MCMCglmm package. We used weakly informative default proper priors for fixed effects and assumed a Gaussian error distribution. For the residual variance, we used an inverse Wishart distribution prior with a low belief parameter nu = 0.002. To estimate inter-individual variances and co-variances, we used parameter-expanded proper priors^50^. We confirmed that results were not affected by changes in the prior specification. For each model, we fitted 5 chains using at least 600 000 iterations, 100 000 burnin and a thinning interval of 100. Chain mixing and convergence were first inspected visually. To confirm model convergence, we ensured that the Gelman-Rubin potential scale reduction factor statistic, calculated with the *gelman.diag* function from the R package coda, was close to 1. The R code with the prior and model specifications is available in the Supplementary data file 3. The estimates of repeatability and inter-individual correlations and their 95% credible intervals were derived from posterior (co)variance distributions using a mode of the posterior density with the function *posterior.mode* and the parameter *adjust* set to 1, and the function *HPDinterval* from the MCMCglmm package, respectively. Because variance must have a positive value, significant repeatability is indicated by confidence intervals not abutting zero, while confidence intervals for significant correlations are not including zero. Full results are reported in Supplemental tables 1-3.

### Reduced representation bisulfite sequencing

Genomic DNA was isolated from micro-dissected dentate gyrus tissue using the QIAamp DNA Micro Kit (QIAGEN) following the manufacturer’s manuals. Libraries for reduced representation bisulfite sequencing (RRBS) were prepared from 100 ng DNA per sample using the Premium RRBS Kit (Diagenode). Final libraries were purified twice using Agencourt AMPure XP beads (Beckman Coulter; 1x bead volume). Quality and concentration of RRBS libraries were determined using the High Sensitivity NGS Fragment Analysis Kit (Advanced Analytical) and a fragment analyzer with capillary size of 33 cm (Advanced Analytical). Sequencing was performed using a NextSeq 5000 platform in a 75 bp single end mode with a minimum sequencing depth of 15 million reads per sample.

### Bioinformatic data analysis of DNA methylation data

Fastq reads were trimmed using Trim Galore 0.4.4 and the function *Cutadapt* 1.8.1 in RRBS mode and mapped against mm10 using Bismark 0.19.0^51^. Detection of differentially methylated cytosines was performed using methylKit v1.5.2^52^. Briefly, methylation levels were extracted from sorted Binary Alignment Map files using the function *processBismarkAln*. Data was filtered for cytosines with a minimum coverage of ten reads and a maximum coverage of 99.9 % percentile in at least three samples per group using the functions *filterByCoverage* and *unite*. Differentially methylated cytosines were identified using the *methDiff* function applying the chi-squared test with multiple testing correction using the Sliding Linear Model (SLIM) method, a significance threshold of *q* < 0.001 and a threshold for absolute cytosine methylation differences higher than 25 %. Overlaps of cytosines was performed using the function *subsetByOverlaps* of the R package Genomic Ranges 3.7^53^. Venn diagrams were generated using the function *draw.pairwise.venn* of the VennDiagram package. Violin plots were created using the R package ggplot2.

Differentially methylated cytosines were annotated to the gene with the nearest transcription start site using data tables downloaded from Ensembl BioMart (download as of 28.05.2019)^54^. Functional gene enrichment analysis was performed using the R packages TopGO and ReactomePA^55^. Gene ontology and pathway enrichment analyses were performed with differentially methylated genes as query lists and all genes covered by RRBS as background lists.

### Data availability statement

Sequencing data have been deposited at GEO (accession number GSE139051). Code for bioinformatic analysis is available upon request.

Data from behavioral analysis and adult neurogenesis have been deposited at Dryad.

## Acknowledgments

This study was financed from basic institutional funds (Helmholtz Association and TU Dresden) and partly supported by the EnergI project funded by Bundesministerium für Bildung und Forschung (BMBF). SZ was supported by a fellowship from the International Max Planck Research School on the Life Course, Berlin. JBL was supported by Coordination for the Improvement of Higher Education Personnel (CAPES), Brazil. The authors thank all members of the Kempermann laboratory for assistance in the logistics of the experiment, Nicole Rund for help with CldU immunohistochemistry and the DRESDEN-concept Genome Center for excellent sequencing service.

## Author contributions

Conceptualization – SZ, ANG, VSA, GK; Methodology – SZ, ANG, SS, RWO, GK; Animal experiment – SS, SZ, JBL, SG; Formal analysis and Visualization – SZ, ANG, SS; Data Curation – SZ, ANG; Writing/ Original Draft – SZ, GK; Writing/ Review and Editing – SZ, ANG, RWO, VSA, JBL, GK; Supervision – SZ, GK.

## References

1. Pennisi, E. The power of personality. Science. 352, 644–647 (2016).

2. Bach, J.-F. The biological individual – The respective contributions of genetics, environment and chance. C. R. Biol. 332, 1065–1068 (2009).

3. Honegger, K. & de Bivort, B. Stochasticity, individuality and behavior. Curr. Biol. 28, R8–R12 (2018).

4. Vogt, G. Stochastic developmental variation, an epigenetic source of phenotypic diversity with far-reaching biological consequences. Journal of Biosciences 40, (2015).

5. Cabeza, R. et al. Maintenance, reserve and compensation: the cognitive neuroscience of healthy ageing. Nat. Rev. Neurosci. 19, 701–710 (2018).

6. Stern, Y. et al. Whitepaper: Defining and investigating cognitive reserve, brain reserve, and brain maintenance. Alzheimer’s Dement. 1–7 (2018).

7. Tang, A. C., Akers, K. G., Reeb, B. C., Romeo, R. D. & McEwen, B. S. Programming social, cognitive, and neuroendocrine development by early exposure to novelty. Proc. Natl. Acad. Sci. U. S. A. 103, 15716–21 (2006).

8. Nithianantharajah, J. & Hannan, A. J. Enriched environments, experience-dependent plasticity and disorders of the nervous system. Nat. Rev. Neurosci. 7, 697–709 (2006).

9. Freund, J. et al. Emergence of individuality in genetically identical mice. Science 340, 756–9 (2013).

10. Freund, J. et al. Association between exploratory activity and social individuality in genetically identical mice living in the same enriched environment. Neuroscience 309, 140–152 (2015).

11. Körholz, J. C. et al. Selective increases in inter-individual variability in response to environmental enrichment in female mice. Elife 7, (2018).

12. Turkheimer, E. & Waldron, M. Nonshared Environment: A Theoretical, Methodological, and Quantitative Review. 126, 78–108 (2000).

13. Kempermann, G. Environmental enrichment, new neurons and the neurobiology of individuality. Nat. Rev. Neurosci. 20, 235–245 (2019).

14. Van Praag, H., Kempermann, G. & Gage, F. H. Neural Consequences of Environmental Enrichment. Nat. Rev. Neurosci. 1, 191–198 (2000).

15. Benito, E. et al. RNA-Dependent Intergenerational Inheritance of Enhanced Synaptic Plasticity after Environmental Enrichment. Cell Rep. 23, 546–554 (2018).

16. Garthe, A., Roeder, I. & Kempermann, G. Mice in an enriched environment learn more flexibly because of adult hippocampal neurogenesis. Hippocampus 26, 261–271 (2016).

17. Kempermann, G., Kuhn, H. G. & Gage, F. H. More hippocampal neurons in adult mice living in an enriched environment. Nature 386, 493–495 (1997).

18. Bennett, E. L., Rosenzweig, M. R., Diamond, M. C., Morimoto, H. & Hebert, M. Effects of Successive Environments on Brain Measures. Physiol. Behav. 12, 621–631 (1974).

19. Bell, A. M., Hankison, S. J. & Laskowski, K. L. The repeatability of behaviour: a meta-analysis. Anim. Behav. 77, 771–783 (2009).

20. Dingemanse, N. J. & Dochtermann, N. A. Quantifying individual variation in behaviour: mixed-effect modelling approaches. J. Anim. Ecol. 82, 39–54 (2013).

21. Kundakovic, M. & Champagne, F. A. Early-Life Experience, Epigenetics, and the Developing Brain. Neuropsychopharmacology 40, 141–153 (2015).

22. Kaas, G. A. et al. TET1 Controls CNS 5-Methylcytosine Hydroxylation, Active DNA Demethylation, Gene Transcription, and Memory Formation. Neuron 79, 1086–1093 (2013).

23. Feng, J. et al. Dnmt1 and Dnmt3a maintain DNA methylation and regulate synaptic function in adult forebrain neurons. Nat. Neurosci. 13, 423–430 (2010).

24. Meissner, A. et al. Reduced representation bisulfite sequencing for comparative high-resolution DNA methylation analysis. Nucleic Acids Res. 33, 5868–5877 (2005).

25. Boyle, P. et al. Gel-free multiplexed reduced representation bisulfite sequencing for large-scale DNA methylation profiling. Genome Biol. 13, R92 (2012).

26. Grunwald, I. C. et al. Kinase-Independent Requirement of EphB2 Receptors in Hippocampal Synaptic Plasticity. Neuron 32, 1027–1040 (2001).

27. Ashton, R. S. et al. Astrocytes regulate adult hippocampal neurogenesis through ephrin-B signaling. Nat. Neurosci. 15, 1399–1406 (2012).

28. Zhang, R.-X. et al. EphB2 in the Medial Prefrontal Cortex Regulates Vulnerability to Stress. Neuropsychopharmacology 41, 2541–2556 (2016).

29. Zhang, J. et al. Increased EphA4-ephexin1 signaling in the medial prefrontal cortex plays a role in depression-like phenotype. Sci. Rep. 7, 7133 (2017).

30. Sheleg, M. et al. Decreased maternal behavior and anxiety in ephrin-A5 ^−/−^ mice.*Genes*, Brain Behav. 16, 271–284 (2017).

31. Dall, S. R. X., Bell, A. M., Bolnick, D. I. & Ratnieks, F. L. W. An evolutionary ecology of individual differences. Ecol. Lett. 15, 1189–1198 (2012).

32. Dingemanse, N. J., Kazem, A. J. N., Réale, D. & Wright, J. Behavioural reaction norms: animal personality meets individual plasticity. Trends Ecol. Evol. 25, 81–89 (2010).

33. Vogt, G. Facilitation of environmental adaptation and evolution by epigenetic phenotype variation: insights from clonal, invasive, polyploid, and domesticated animals. Environ. Epigenetics 3, 1–17 (2017).

34. Garthe, A., Huang, Z., Kaczmarek, L., Filipkowski, R. K. & Kempermann, G. Not all water mazes are created equal: Cyclin D2 knockout mice with constitutively suppressed adult hippocampal neurogenesis do show specific spatial learning deficits. *Genes*, Brain Behav. 13, 357–364 (2014).

35. Sahay, A. et al. Increasing adult hippocampal neurogenesis is sufficient to improve pattern separation. Nature 472, 466–470 (2011).

36. David, D. J. et al. Neurogenesis-Dependent and -Independent Effects of Fluoxetine in an Animal Model of Anxiety/Depression. Neuron 62, 479–493 (2009).

37. Anacker, C. & Hen, R. Adult hippocampal neurogenesis and cognitive flexibility - linking memory and mood. Nat. Rev. Neurosci. 18, 335–346 (2017).

38. Busche, S. et al. Population whole-genome bisulfite sequencing across two tissues highlights the environment as the principal source of human methylome variation. Genome Biol. 16, 290 (2015).

39. Gunasekara, C. J. et al. A genomic atlas of systemic interindividual epigenetic variation in humans. Genome Biol. 20, 105 (2019).

40. Weaver, I. C. G. et al. Epigenetic programming by maternal behavior. Nat. Neurosci. 7, 847–54 (2004).

41. Fischer, A., Sananbenesi, F., Wang, X., Dobbin, M. & Tsai, L.-H. Recovery of learning and memory is associated with chromatin remodelling. Nature 447, 178–183 (2007).

42. Muotri, A. R., Zhao, C., Marchetto, M. C. N. & Gage, F. H. Environmental influence on L1 retrotransposons in the adult hippocampus. Hippocampus 19, 1002–1007 (2009).

43. Choi, C., Kim, T., Chang, K. T. & Min, K. DSCR 1-mediated TET 1 splicing regulates miR-124 expression to control adult hippocampal neurogenesis. EMBO J. 38, (2019).

44. Muotri, A. R. et al. L1 retrotransposition in neurons is modulated by MeCP2. Nature 468, 443–446 (2010).

45. Campbell, R. R. & Wood, M. A. How the epigenome integrates information and reshapes the synapse. Nat. Rev. Neurosci. 20, 133–147 (2019).

46. Noguchi, K., Gel, Y. R., Brunner, E. & Konietschke, F. nparLD: An *R* Software Package for the Nonparametric Analysis of Longitudinal Data in Factorial Experiments. J. Stat. Softw. 50, 1–23 (2012).

47. Wickham, H. ggplot2. Wiley Interdiscip. Rev. Comput. Stat. 3, 180–185 (2011).

48. Nakagawa, S. & Schielzeth, H. Repeatability for Gaussian and non-Gaussian data: a practical guide for biologists. Biol. Rev. 85, no-no (2010).

49. Hadfield, J. D. MCMC Methods for Multi-Response Generalized Linear Mixed Models: The MCMCglmm *R* Package. J. Stat. Softw. 33, 1–22 (2010).

50. Gelman, A. Prior distributions for variance parameters in hierarchical models (comment on article by Browne and Draper). Bayesian Anal. 1, 515–534 (2006).

51. Krueger, F. & Andrews, S. R. Bismark: a flexible aligner and methylation caller for Bisulfite-Seq applications. Bioinformatics 27, 1571–1572 (2011).

52. Akalin, A. et al. methylKit: a comprehensive R package for the analysis of genome-wide DNA methylation profiles. Genome Biol. 13, R87 (2012).

53. Lawrence, M., et al. Software for Computing and Annotating Genomic Ranges. PLOS Comput. Biol. 9, 1–10 (2013).

54. Zerbino, D. R., et al. Ensembl 2018. Nucleic Acid Res. 46, 754–761 (2018).

55. Yu, G. & He, Q. Molecular BioSystems ReactomePA: an R / Bioconductor package for reactome pathway analysis and visualization. Mol. Biosyst. 12, 477–479 (2016).

