## Supplementary data for "Early-life environmental enrichment generates persistent individualized behavior in mice"

Sara Zocher*, Susan Schilling*, Anna N. Grzyb*, Vijay S. Adusumilli, Jadna Bogado Lopes, Sandra Günther, Rupert W. Overall, Gerd Kempermann^#^

German Center for Neurodegenerative Diseases (DZNE) Dresden, Tatzberg 41, 01307 Dresden, Germany

Center for Regenerative Therapies Dresden (CRTD), Technische Universität Dresden, Fetscherstraße 105, 01307 Dresden, Germany

*These authors contributed equally.

^#^ Corresponding author.

Content of Supplementary Information:

Supplementary Figures 1-4 with figure legends

Supplementary Tables 1-3

Captions for Supplementary Data files 1-3

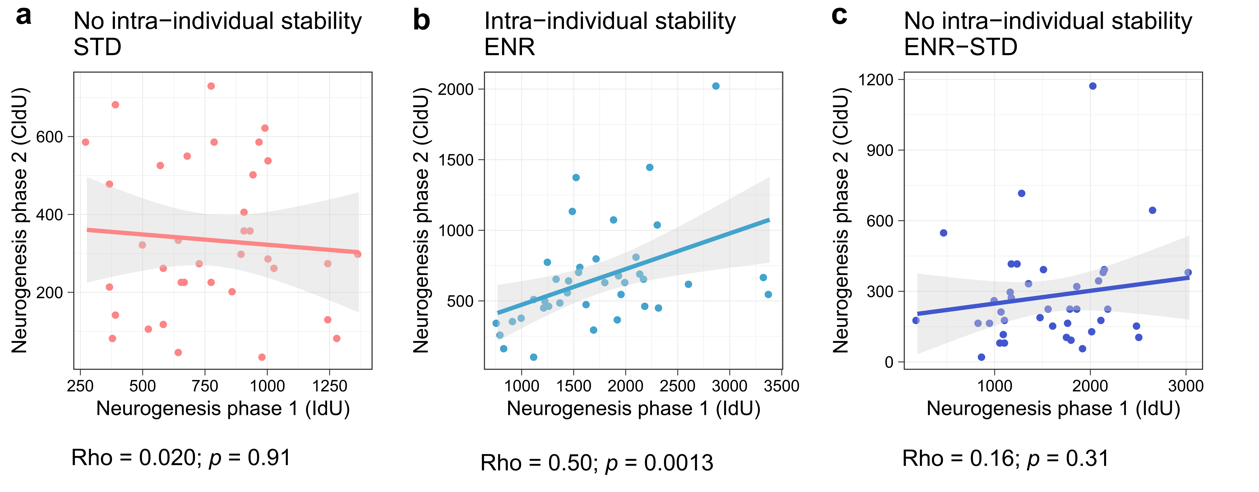

**Supplemental Fig. 1:** ENR stabilized levels of adult hippocampal neurogenesis during aging. **a-c,** show correlations of individual levels of adult hippocampal neurogenesis between phase 1 and phase 2 (IdU vs CldU) in mice separated by groups. Depicted are Spearman’s rho and *p*-value of the rank correlation.

Figure refers to content in Figure 3.

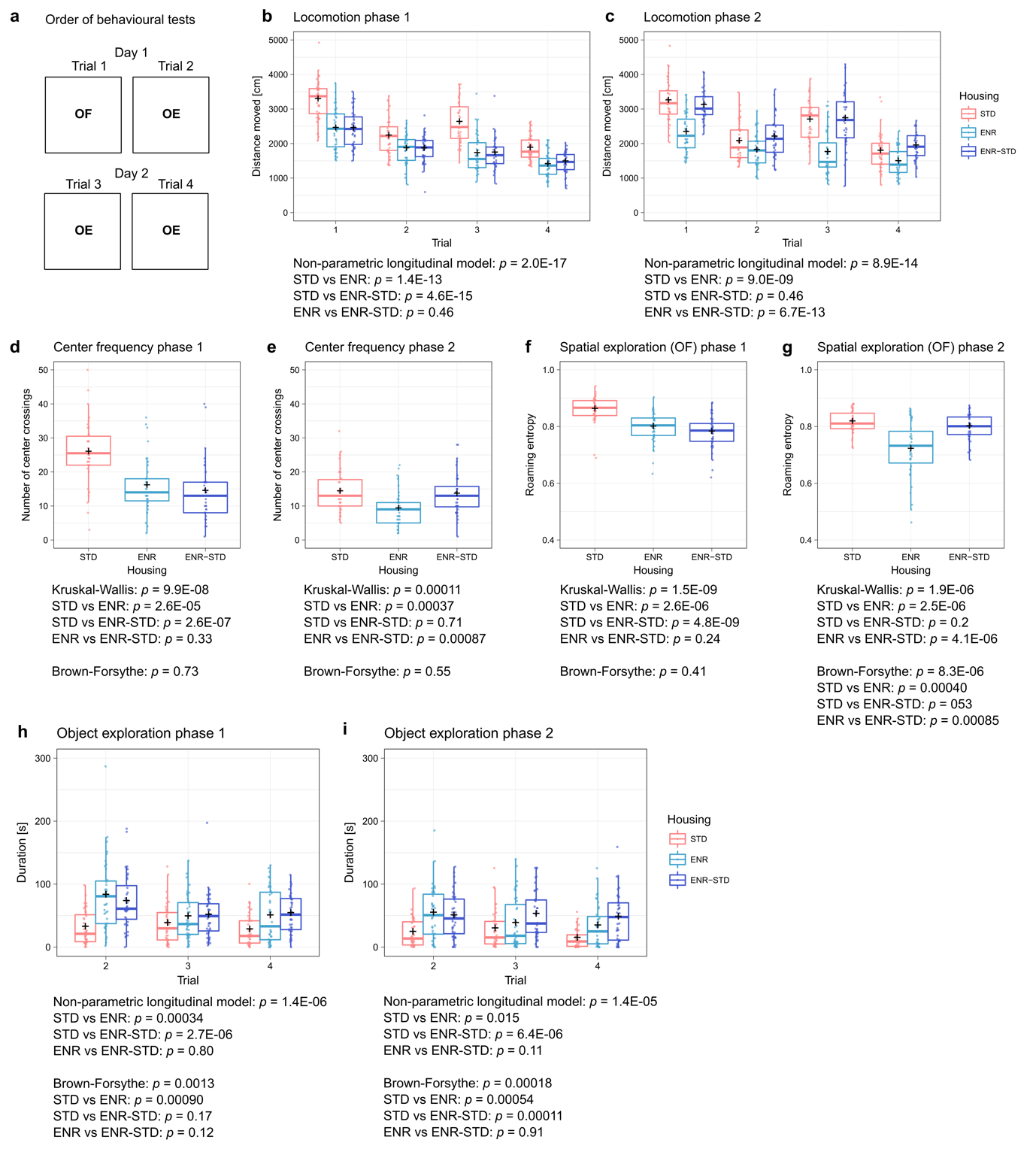

**Supplemental Fig. 2:** Results of cross-sectional behavioral testing. ENR-induced behavioral changes in the open field are plastic towards environmental stimulation and reversed after withdrawal from ENR, while ENR-induced changes in object exploration are maintained after environmental change. **a,** Timeline for behavioral testing in phase 1 and phase 2. One trial of open field (OF) test and three trials of object exploration (OE) test were performed in the same arena on two consecutive days. **b,** Locomotion during individual trials of OF and OE tests. At the end of phase 1, ENR and ENR-STD mice traveled shorter distances compared to STD mice. **c,** At the end of phase 2, ENR mice maintained reduced levels of locomotion in OF and OE tests, while ENR-STD mice traveled distances similar to STD mice. **d,** ENR increased the frequency of center passage in OF test. **e,** Increase in frequency of center passage was not maintained in ENR-STD after withdrawal. **f,** ENR mice showed reduced roaming entropy in the open field compared to STD mice. **g,** Withdrawal from ENR increased roaming entropy in the open field to the levels observed in STD mice, while ENR mice show reduced mean and higher variance of the values. **h-i,** Object exploration during all three trials of the object exploration test in phase 1 and phase 2. See Supplemental data 1 for detailed statistical results.

Figure refers to content in Figure 4.

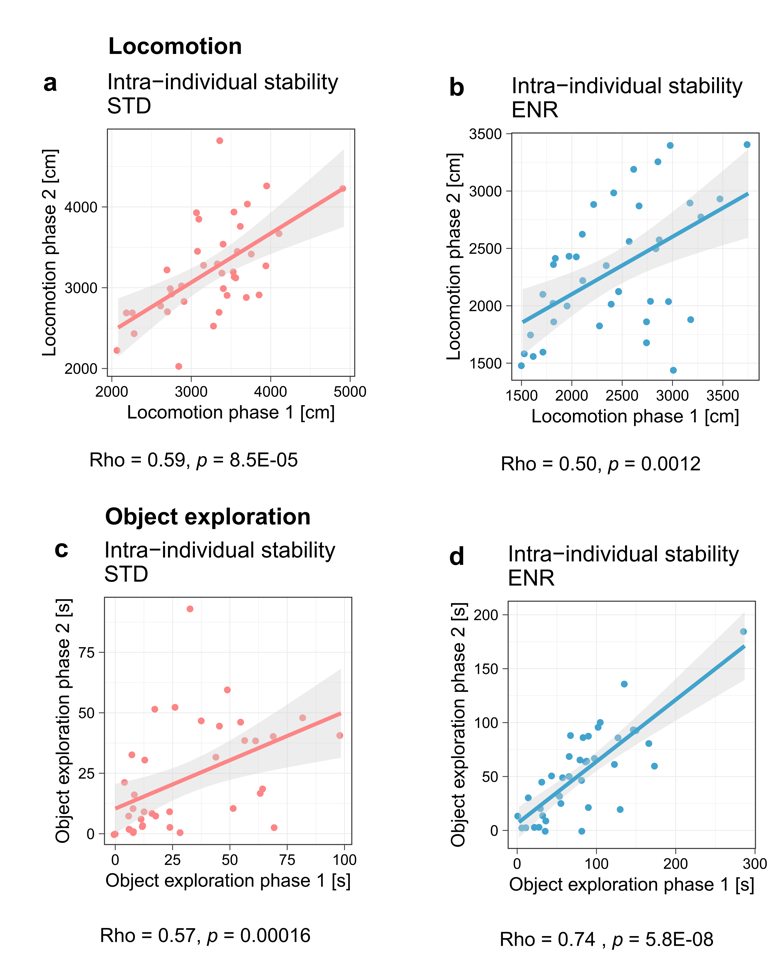

**Supplemental Fig. 3:** STD and ENR mice show intra-individual stability in object exploration and locomotion. **a-b,** Correlations of individual locomotion in open field test between phase 1 and phase 2. **c-d,** Correlations of object exploration (time around objects) in the first trial of the object exploration test between phase 1 and phase 2. Depicted are Spearman’s rho and *p*-value of the rank correlation.

Figure refers to content in Figure 4.

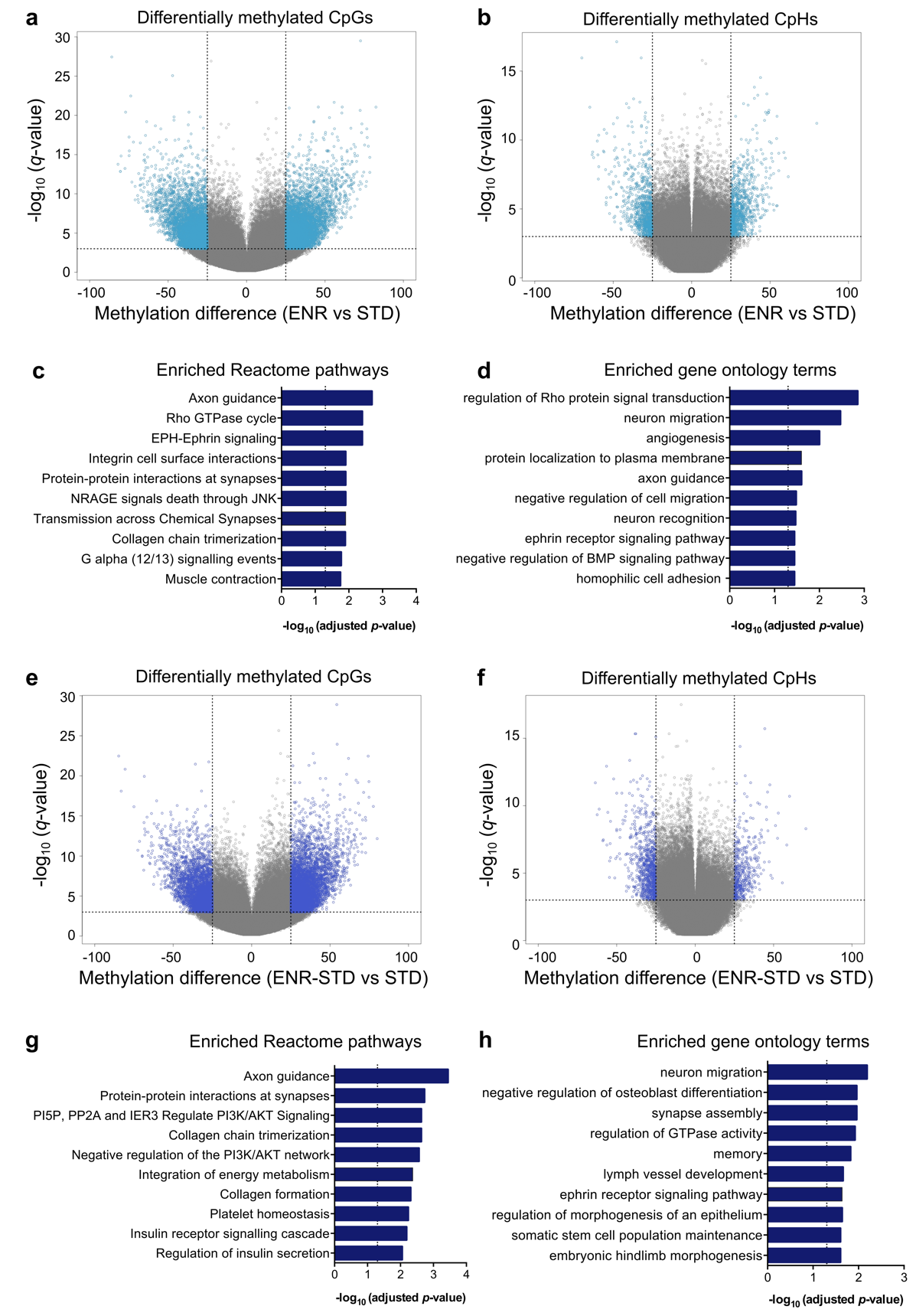

**Supplemental Fig. 4:** **a-b,** Volcano plot depicting significantly differentially methylated CpGs and CpHs (*q* < 0.001, methylation difference > 25 %) between ENR and STD mice in pale blue. **c-d,** Genes with ENR-induced differentially methylated cytosines (CpGs or CpHs) are enriched at neuronal plasticity pathways. **e-f,** Significantly differentially methylated cytosines between ENR-STD and STD mice depicted in dark blue. **g-h,** Gene ontology and pathway enrichment of genes differentially methylated between ENR-STD and STD.

Figure refers to content in Figure 5.

**Supplementary Table 1:** Variance estimates from linear mixed models with 95 % credibility intervals (CI).

| **Phenotype** | **Housing** | **Vind** | **Vind**  **CI_low** | **Vind**  **CI_up** | **Vres** | **Vres**  **CI_low** | **Vres**  **CI_up** | **R** | **R**  **CI_low** | **R**  **CI_up** |
| --- | --- | --- | --- | --- | --- | --- | --- | --- | --- | --- |
| RE T1 | ENR | 0.00 | 0.00 | 0.14 | 0.97 | 0.81 | 1.20 | 0.00 | 0.00 | 0.13 |
| RE T3 | ENR | 0.10 | 0.05 | 0.19 | 0.52 | 0.44 | 0.59 | 0.17 | 0.09 | 0.28 |
| RE T4 | ENR | 0.13 | 0.07 | 0.24 | 0.66 | 0.60 | 0.74 | 0.17 | 0.09 | 0.27 |
| RE T5 | ENR | 0.20 | 0.12 | 0.34 | 0.63 | 0.56 | 0.71 | 0.24 | 0.16 | 0.36 |
| RE T6 | ENR | 0.27 | 0.16 | 0.48 | 0.83 | 0.74 | 0.92 | 0.25 | 0.17 | 0.37 |
| RE T7 | ENR | 0.12 | 0.07 | 0.23 | 0.75 | 0.67 | 0.84 | 0.14 | 0.09 | 0.24 |
| RE T8 | ENR | 0.23 | 0.13 | 0.42 | 0.80 | 0.72 | 0.92 | 0.23 | 0.14 | 0.34 |
| Neurogenesis | STD | 0.00 | 0.00 | 0.03 | 0.11 | 0.08 | 0.16 | 0.01 | 0.00 | 0.23 |
| Neurogenesis | ENR | 0.17 | 0.00 | 0.40 | 0.32 | 0.20 | 0.54 | 0.38 | 0.04 | 0.63 |
| Neurogenesis | ENR-STD | 0.01 | 0.00 | 0.16 | 0.36 | 0.24 | 0.52 | 0.02 | 0.00 | 0.34 |
| Locomotion | STD | 15.07 | 7.35 | 29.55 | 9.84 | 6.45 | 16.29 | 0.64 | 0.39 | 0.79 |
| Locomotion | ENR | 16.35 | 5.32 | 33.12 | 15.11 | 9.57 | 25.58 | 0.55 | 0.26 | 0.74 |
| Locomotion | ENR-STD | 0.52 | 0.00 | 9.36 | 17.01 | 11.07 | 25.15 | 0.02 | 0.00 | 0.39 |
| Initial object exploration | STD | 3.38 | 1.52 | 6.67 | 2.40 | 1.53 | 4.03 | 0.60 | 0.35 | 0.78 |
| Initial object exploration | ENR | 7.23 | 4.13 | 12.69 | 2.71 | 1.74 | 4.36 | 0.75 | 0.56 | 0.86 |
| Initial object exploration | ENR-STD | 4.03 | 1.56 | 7.96 | 3.61 | 2.23 | 5.78 | 0.59 | 0.29 | 0.74 |

Vind – inter-individual variance component; Vres – residual variance component; R – repeatability; CI_low – lower confidence interval; CI_up – upper confidence interval.

**Supplementary Table 2:** Comparison of models with heterogeneous inter-individual and residual variances with models assuming homogeneous variance. Reported are values of deviance information criterion (DIC) for the full and simplified models and the difference between simplified and full models. Reduction of ∆DIC by at least 2 units can suggests better fit to the data.

| **Phenotype** | **Full model** | **Homogeneous Vind** | **∆DIC (full – simpler model)** | **Homogeneous Vres** | **∆DIC (full – simpler model)** |
| --- | --- | --- | --- | --- | --- |
| Neurogenesis | 373.5 | 387.1 | 13.6 | 380.1 | 6.6 |
| RE | 9591.0 | 9676.3 | 36.8 | 9627.7 | 36.8 |
| RE (time blocks T3-T8) | 8902.6 | 8957.3 | 54.7 | 8934.5 | 31.9 |
| Locomotion | 1365.1 | 1369.9 | 4.9 | 1371.6 | 6.6 |
| Initial object exploration | 1017.2 | 1016.6 | -0.6 | 1016.8 | -0.4 |

Vind – inter-individual variance component; Vres – residual variance component.

**Supplementary Table 3:** Posterior mode of inter-individual correlation of RE between time blocks with 95 % credibility intervals.

| **Time blocks** | **Pearson’s r** | **CI_low** | **CI_up** |
| --- | --- | --- | --- |
| T3:T1 | 0.15 | -0.78 | 0.87 |
| T4:T1 | 0.14 | -0.84 | 0.81 |
| T5:T1 | 0.08 | -0.89 | 0.92 |
| T6:T1 | -0.11 | -0.91 | 0.89 |
| T7:T1 | 0.00 | -0.89 | 0.91 |
| T8:T1 | 0.07 | -0.86 | 0.95 |
| T4:T3 | 0.71 | 0.33 | 0.90 |
| T5:T3 | 0.87 | 0.59 | 0.97 |
| T6:T3 | 0.75 | 0.44 | 0.94 |
| T7:T3 | 0.83 | 0.52 | 0.97 |
| T8:T3 | 0.82 | 0.53 | 0.96 |
| T5:T4 | 0.84 | 0.61 | 0.95 |
| T6:T4 | 0.82 | 0.59 | 0.95 |
| T7:T4 | 0.80 | 0.52 | 0.94 |
| T8:T4 | 0.79 | 0.54 | 0.94 |
| T6:T5 | 0.91 | 0.76 | 0.99 |
| T7:T5 | 0.93 | 0.77 | 0.99 |
| T8:T5 | 0.93 | 0.77 | 0.99 |
| T7:T6 | 0.95 | 0.79 | 0.99 |
| T8:T6 | 0.93 | 0.79 | 0.99 |
| T8:T7 | 0.93 | 0.75 | 0.99 |

CI_low – lower confidence interval; CI_up – upper confidence interval.

Captions for Supplementary Data 1-3

**Supplementary Data 1**

Details of statistical analysis of behavioral data and adult hippocampal neurogenesis. Data related to Fig. 3-4 and Suppl. Fig. 2.

**Supplementary Data 2**

Lists of differentially methylated cytosines between ENR and STD mice (sheet 1), differentially methylated cytosines between ENR-STD and STD (sheet 2) and list of ENR-induced differentially methylated cytosines maintained after withdrawal (sheet 3). Data related to Fig. 5 and Suppl. Fig. 4.

**Supplementary Data 3**

R code for mixed linear models and repeatability calculation related to results in Fig. 2-4.
